# Macrophages only sense infectious SARS-CoV-2 when they express sufficient ACE2 to permit viral entry, where rapid cytokine responses then limit viral replication

**DOI:** 10.1101/2022.03.22.485248

**Authors:** Larisa I Labzin, Keng Yih Chew, Kathrin Eschke, Xiaohui Wang, Tyron Esposito, Claudia J Stocks, James Rae, Ralph Patrick, Helen Mostafavi, Brittany Hill, Teodor E. Yordanov, Caroline L Holley, Stefan Emming, Svenja Fritzlar, Francesca L. Mordant, Daniel P. Steinfort, Kanta Subbarao, Christian M. Nefzger, Anne K Lagendijk, Emma Gordon, Robert Parton, Kirsty R. Short, Sarah L. Londrigan, Kate Schroder

## Abstract

Macrophages are key cellular contributors to COVID-19 pathogenesis. Whether SARS-CoV-2 can enter macrophages, replicate and release new viral progeny remains controversial. Similarly, whether macrophages need to sense replicating virus to drive cytokine release is also unclear. Macrophages are heterogeneous cells poised to respond to their local microenvironment, and accordingly, the SARS-CoV-2 entry receptor ACE2 is only present on a subset of macrophages at sites of human infection. Here, we use in vitro approaches to investigate how SARS-CoV-2 interacts with ACE2-negative and ACE2-positive human macrophages and determine how these macrophage populations sense and respond to SARS-CoV-2. We show that SARS-CoV-2 does not replicate within ACE2-negative human macrophages and does not induce pro-inflammatory cytokine expression. By contrast, ACE2 expression in human macrophages permits SARS-CoV-2 entry, replication, and virion release. ACE2-expressing macrophages sense replicating virus to trigger pro-inflammatory and anti-viral programs that limit virus release. These combined findings resolve several controversies regarding macrophage-SARS-CoV-2 interactions and identify a signaling circuit by which macrophages sense SARS-CoV-2 cell entry and respond by restricting viral replication.

**One sentence summary:** Lack of macrophage ACE2 expression precludes SARS-CoV-2 entry and sensing, while ACE2-expressing macrophages sense intramacrophage SARS-CoV-2 replication to induce rapid anti-viral responses that limit new virion release.

## INTRODUCTION

Severe Acute Respiratory Syndrome Coronavirus 2 (SARS-CoV-2), belonging to the *Coronaviridae* family, is an enveloped, single-stranded RNA virus with a positive-sense genome. Many human respiratory viruses, including SARS-CoV-2, infect epithelial cells lining the upper and lower airways, resulting in productive replication and the release of newly synthesized infectious viral particles. The virus Spike (S) glycoprotein facilitates entry into target epithelial cells by binding to the surface-expressed angiotensin-converting enzyme 2 (ACE2) receptor (*1*). The well-characterized ACE2-Spike interaction exposes a critical S cleavage site (S2) that can be cleaved by the host serine protease TMPRSS2 (transmembrane protease serine 2), also expressed on the plasma membrane. This allows the fusion of the viral and cellular membranes, followed by the release of viral RNA directly into the cytoplasm (reviewed in (*2*) and (*3*)). SARS-CoV-2, particularly omicron variants, can also attach to ACE2 and enter cells via the endocytic pathway. Here, the S protein is cleaved by endosomal proteases to allow fusion between the viral and endosomal membranes (*4*). Host cell ribosomes immediately translate the infecting, positive sense RNA genome into two large polyproteins. These polyproteins are proteolytically processed to generate individual viral proteins required for replication. New virions are assembled in the endoplasmic reticulum and Golgi of the host cell, before secretion from the cell via exocytosis or through a lysosomal egress pathway (*3*). Our understanding of the SARS-CoV-2 replication cycle comes from studies in primary epithelial cells or epithelial cell lines. How this may differ in other potentially susceptible cell types, including distinct macrophage populations, remains unclear.

Effective host defense against infection relies on accurate and timely immune detection. Accumulating evidence suggests that severe COVID-19 results from a failure of early host-interferon signaling to control SARS-CoV-2, followed by exacerbated pro-inflammatory responses driving tissue damage. Airway epithelial cells, the primary target for SARS-CoV-2 infection (*5*), respond by releasing anti-viral and pro-inflammatory cytokines (*6, 7*). Airway-resident or newly recruited macrophages also appear to be a key source of pro-inflammatory cytokines in severe COVID-19 (*8, 9*), with macrophage-derived cytokines implicated in the severe pathology observed in patients (*10–13*).

Macrophages are sentinel innate immune cells that defend the airways during respiratory viral infection. Yet, and in contrast to epithelial cells, infection of human macrophages results in an abortive replication cycle for many respiratory viruses (including seasonal influenza A viruses (*14*) and rhinovirus (*15*)), despite macrophages supporting the early stages of infection (entry and synthesis of new viral RNA and protein). Therefore, macrophages can function as a viral ‘dead end’ to limit viral dissemination (*16*) and can sense infectious viral particles, infected cells, and tissue damage to orchestrate anti-viral and pro-inflammatory programs (*17–19*). Single-cell RNA sequencing has revealed viral RNA within macrophage populations in human COVID-19 lungs (*8, 20*) and *ex vivo* lung explants (*21*). Similarly, immunostaining of COVID-19 autopsy lungs showed macrophages containing viral antigens (Spike and the RNA-dependent RNA polymerase: RdRp) *in situ* (*22*). Whether SARS-CoV-2 enters these macrophages via ACE2 remains unclear, as is the fate of intramacrophage SARS-CoV-2. These in vivo studies did not ascertain if SARS-CoV-2 productively replicates within macrophages to produce new viral RNA and protein leading to new virion assembly, nor whether macrophages release newly synthesized infectious virions. Further, previous studies did not elucidate whether SARS-CoV-2 needs to enter and actively replicate in macrophages to trigger macrophage anti-viral and pro-inflammatory responses.

In vitro models of SARS-CoV-2 interactions with macrophages have focused on human monocyte-derived macrophages (HMDM). While there is consensus that HMDMs exposed to SARS-CoV-2 do not release newly-synthesized infectious virions (*7, 12, 13, 23–25*), some studies report that HMDMs support the early stages of infection that include viral entry and replication (i.e., viral RNA replication and protein synthesis) (*12, 13, 23*). In contrast, others report that HMDM are refractory to SARS-CoV-2 entry (*7, 25*). Accordingly, whether SARS-CoV-2 exposure induces HMDM pro-inflammatory and anti-viral signaling or whether this requires active viral entry and replication is also controversial. Multiple reports suggest that macrophage challenge with SARS-CoV2 triggers pro-inflammatory responses (*12, 13, 26*). In contrast, others report that SARS-CoV-2 challenge does not activate macrophage inflammatory functions, consistent with a lack of viral entry and the early stages of viral replication (*7, 25*).

The current study investigates whether macrophage ACE2 expression dictates macrophage susceptibility to SARS-CoV-2 entry and replication and how this shapes macrophage inflammatory responses. We found that SARS-CoV-2 cannot enter or replicate within ACE2-negative macrophages, and these cells do not produce pro-inflammatory cytokines in response to viral challenge. In contrast, macrophages engineered to ectopically express ACE2 support SARS-CoV-2 entry, replication, and new virion release. ACE2-expressing macrophages sense newly synthesized viral RNA and induce pro-inflammatory and anti-viral mediator expression. This rapid induction of anti-viral programs in ACE2-expressing macrophages limits ongoing virion release. Thus, we identify two blocks to SARS-CoV-2 infection and replication in ACE2-negative and -positive macrophages. ACE2-negative macrophages are refractory to SARS-CoV-2 infectious entry and do not support the early stages of replication. In contrast, ACE2-positive macrophages are susceptible to the early stages of infection and replication but mount a second post-entry block that constrains the production and release of newly synthesized infectious virions.

## RESULTS

### In vitro models of ACE2-positive and ACE2-negative macrophages

Given that ACE2 is the primary receptor for SARS-CoV-2, we anticipated that ACE2 expression would determine the susceptibility and permissivity of macrophages to SARS-CoV-2 infection and replication. We reanalyzed published scRNA-seq lung data from three studies (*20, 27, 28*), representing two sets of control and two sets of COVID-19-infected individuals. We aligned the datasets to a common cell type reference to study *ACE2* mRNA expression amongst myeloid cell types (e.g., infiltrating monocytes, monocyte-derived macrophages, and resident alveolar macrophages). Less than 2% of all cell types expressed *ACE2* mRNA, including alveolar epithelial AT1 and AT2 cells targeted by SARS-CoV-2 (**fig. S1**). In control and COVID-19-infected lungs, a small proportion (<0.05%) of monocyte-derived macrophages expressed *ACE2* mRNA. Very few tissue-resident macrophages and transitioning monocyte-derived macrophages expressed *ACE2* mRNA in control lungs, though this proportion increased in the COVID-19-infected lung. Monocytes did not express *ACE2* mRNA in either condition **(fig. S1)**. These results are consistent with published scRNA sequencing analyses on *ACE2* mRNA expression across cell types (*20, 27–29*). A recent preprint reported ACE2-positive macrophages in lung explants infected *ex vivo*, as determined by immunofluorescence (*30*). These data suggest that both ACE2-positive and ACE2-negative macrophages are present in SARS-CoV-2-infected lungs.

We next assessed ACE2 expression in HMDM cultured from blood monocytes to determine their suitability as an in vitro cell model for ACE2-positive or ACE2-negative macrophages. Monitoring ACE2 mRNA and protein revealed low *ACE2* mRNA expression in HMDM, and undetectable levels of ACE2 protein (**Fig. 1A, B**), even when HMDM were treated with IFNβ (**Fig. 1B**). HMDM thus resemble ACE2-negative macrophages in COVID-19 lungs.

**Figure 1:**
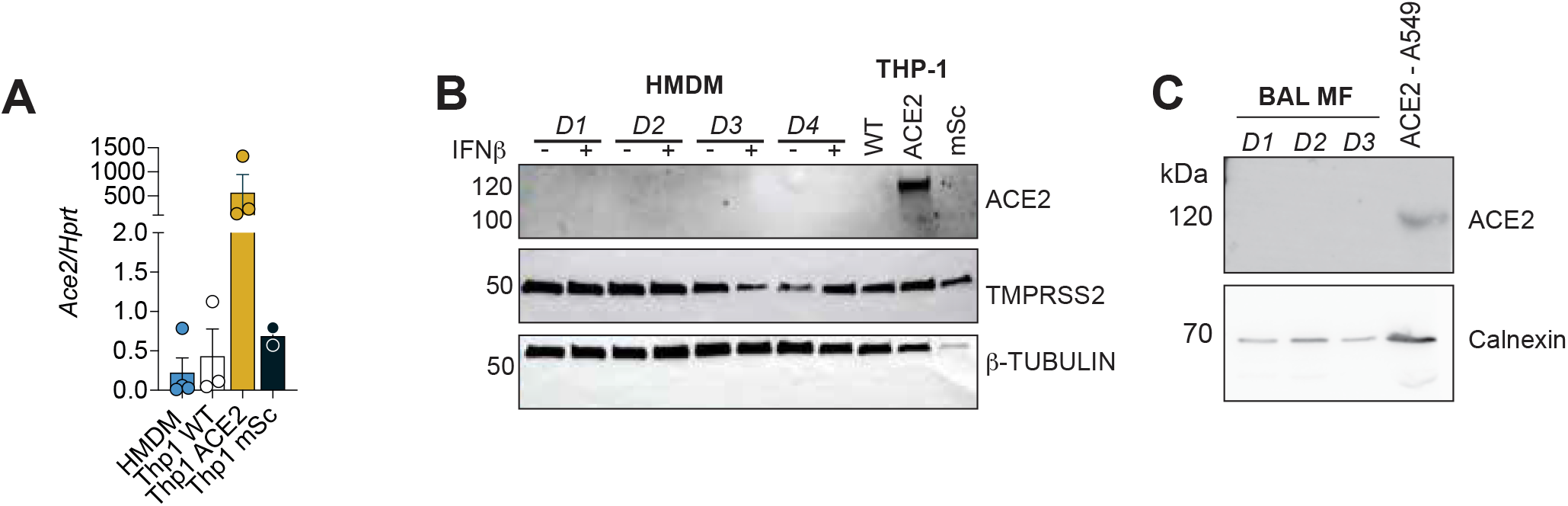
ACE2 is not expressed on HMDM or BAL but can be ectopically expressed in THP-1 cells. A: HMDM and THP-1 cells were analyzed by qPCR for *ACE2* mRNA expression, with each data point showing an independent donor or experiment (n = 3). B: HMDM were stimulated with IFNβ (10 ng/ml) for six h, and protein extracts were analyzed by immunoblot alongside extracts from THP-1 cells (WT, THP-1-ACE2, THP-1-mSc). C: BAL macrophages from 3 donors were adhered overnight and lysed. The expression of ACE2 in BAL macrophages was analyzed by immunoblot relative to loading control (Calnexin). Lysate from A549 cells overexpressing ACE2 was used as a positive control.

The lung-resident macrophage populations are immune sentinels of the airways, with distinct origins and properties to HMDM (*31*). Along with airway epithelial cells, airway macrophages are prime targets for infection by many respiratory viruses. We thus investigated the ACE2 expression status of this lung-resident macrophage population. We utilized airway macrophages from bronchoalveolar lavage (BAL) as representative resident lung macrophages and observed that their ACE2 protein expression was below detection levels (**Fig. 1C**). Together, this data indicates that HMDM and BAL macrophages cultured ex vivo model ACE2-negative macrophage populations in vivo.

To model ACE2-positive cells, we used lentiviral transduction to ectopically express either ACE2 (untagged) or a control protein (mScarlet: mSc) in the THP-1 monocytic cell line prior to their differentiation into macrophage-like cells with phorbol-myristate-acetate (PMA). We confirmed that ectopic ACE2 expression in THP-1 cells yielded readily detectable ACE2 mRNA and protein **(Fig. 1A, 1B)**. The surface protease TMPRSS2, which is required for S protein cleavage, was also readily detected at the protein level in HMDM, THP-1-ACE2, and THP-1-mSc cells **(Fig. 1B)**, indicating its presence in both ACE2-positive and ACE2-negative macrophages.

### ACE2-negative macrophages do not support SARS-CoV-2 entry and replication

Despite the small proportion of ACE2-positive macrophages at sites of human SARS-CoV-2 infection ((*20, 27*), **fig S1**), many macrophages are positive for SARS-CoV-2 RNA or protein in vivo (*8, 20, 22*), potentially reflecting an ACE2-independent mechanism of viral entry into macrophages. We thus investigated whether SARS-CoV-2 can enter HMDMs, which do not express ACE2 (**Fig. 1A, 1B)**. We left the viral inoculum on HMDM to maximize the uptake of SARS-CoV-2 virions. For these experiments, we included SARS-CoV-2-permissive Calu-3 epithelial cells as a positive control for viral infection and replication. Calu-3 cells were susceptible to infection at high and low doses (MOI 5 and 0.5). We detected an increase in cell-associated viral RNA between two hours (indicative of input virus) and 24 h post-infection (p.i.), indicating newly synthesized viral RNA and active replication (**Fig. 2A**). Consistent with productive infection in Calu-3 cells, we also detected viral sub-genomic RNA (indicating new viral RNA synthesis; **Fig. S2A**) and newly synthesized viral nucleoprotein (NP, increased from two to 72 h; **Fig. 2B**). In contrast, cell-associated viral RNA levels did not rise over the same time course in HMDM infected with either MOI dosage (**Fig. 2C**). Viral sub-genomic RNA was barely detectable in HMDM (**Fig. S2B**), and while we could detect NP expression in HMDM at two hours p.i., NP levels were barely detectable 24 h p.i. (**Fig. 2D**). These data indicate that HMDM are not susceptible to SARS-CoV-2 infection and do not support the early stages of viral replication, including the production of newly synthesized viral RNA and protein.

**Figure 2:**
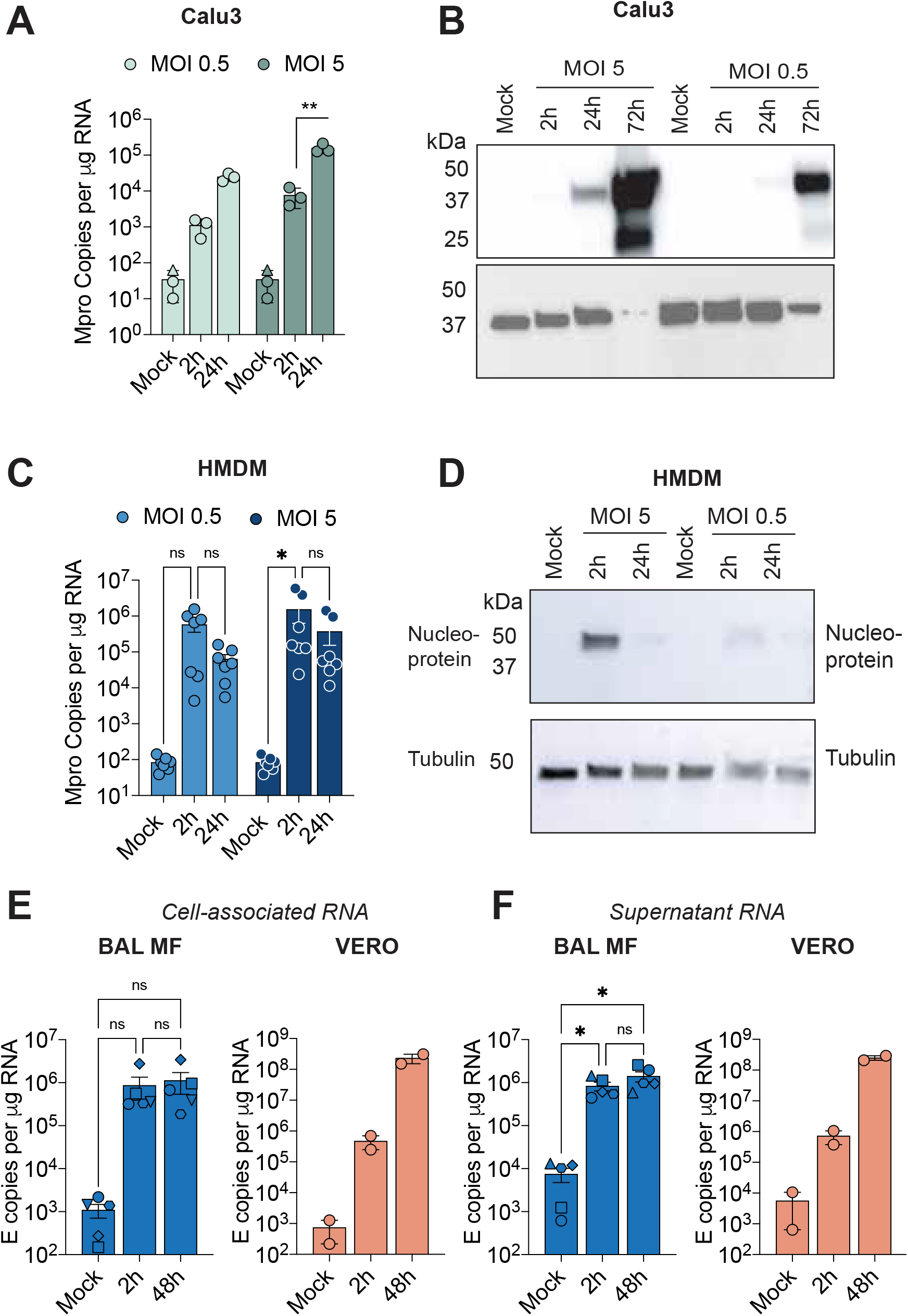
ACE2-negative HMDM and BAL do not support SARS-CoV-2 entry or early-stage replication. A-B: Calu-3 cells (A, B) or HMDM (C,D) were infected as indicated, and the virus was left on the cells. Viral RNA isolated from cells was measured by qPCR (A, C), or viral protein by immunoblot (B,D). Blots are representative of three independent experiments (Calu-3) or three independent donors (HMDM). E-F: BAL macrophages (MOI 1) or Vero cells (MOI 0.5) were infected with SARS-CoV-2 for one h before the virus was removed, and cell-associated viral RNA (E) or viral RNA released in cell-free supernatants (F) was analyzed by qPCR of SARS-CoV-2 E gene at 2- and 48-h post-infection. Data are mean + SEM, with each point representing an individual donor (HMDM, BAL macrophages; n = 5) or independent experiments (Calu-3, VERO; n = 2-3). Asterisks indicate significance: p ≤ 0.05 (*), p ≤ 0.001 (**), p ≤ 0.0001 (***) (two-way ANOVA, Tukey’s multiple comparison test).

We next tested whether BAL macrophages support ACE2-independent viral entry and replication by challenging these cells with SARS-CoV-2 at an MOI of 1. SARS-CoV-2-challenged BAL macrophages showed no increase in cell-associated viral RNA (indicative of the early stages of virus replication) between two and 48 h post-infection, in contrast to the SARS-CoV-2-susceptible control cell line, VERO E6 (**Fig. 2E**). An increase in viral RNA isolated from cell-free VERO E6 supernatants between two and 48 h post-infection indicated productive viral replication. However, we did not observe an increase in viral RNA in supernatants of SARS-CoV-2 infected BAL macrophages (**Fig. 2F**). As for HMDM, these data indicate that ACE2-negative BAL macrophages are not permissive to the early stages of SARS-CoV-2 infection and replication.

### ACE2-positive macrophages support viral entry, replication, and new virion release

We next investigated whether ACE2-positive macrophages could support SARS-CoV-2 entry and replication. We challenged THP-1-ACE2 **(Fig. 1A, 1B)** with SARS-CoV-2 (MOI 0.5 and 5) and compared viral replication and release to THP-1-mSc (ACE2-negative) and Calu-3 control cells. Cell-associated viral RNA significantly increased from 0 h (input) to 24 h p.i. for both MOIs in THP-1-ACE2 cells before plateauing from 24 to 72 h p.i., indicating cell susceptibility and early-stage viral RNA replication (**Fig. 3A, 3B)**. By contrast, viral RNA levels did not increase in the THP-1-mSc control cells at any time (**Fig. 3A, 3B**), similar to our observations in other ACE2-negative macrophages (HMDM and BAL**; Fig. 2C, 2E, 2F)**. Calu-3 cells support viral replication and virion release (*32*); accordingly, SARS-CoV-2 RNA levels continued to rise over time (**Fig. 3A, 3B**). Consistent with new viral protein synthesis, we observed robust NP staining at 24 h post-infection in THP-1-ACE2 but not THP-1-mSc cells (**Fig. 3C**). Only 16 % of THP-1-ACE2 cells were NP positive at 24 h p.i. (**Fig. 3D**), indicating heterogeneity in the cellular response to SARS-CoV-2.

**Figure 3:**
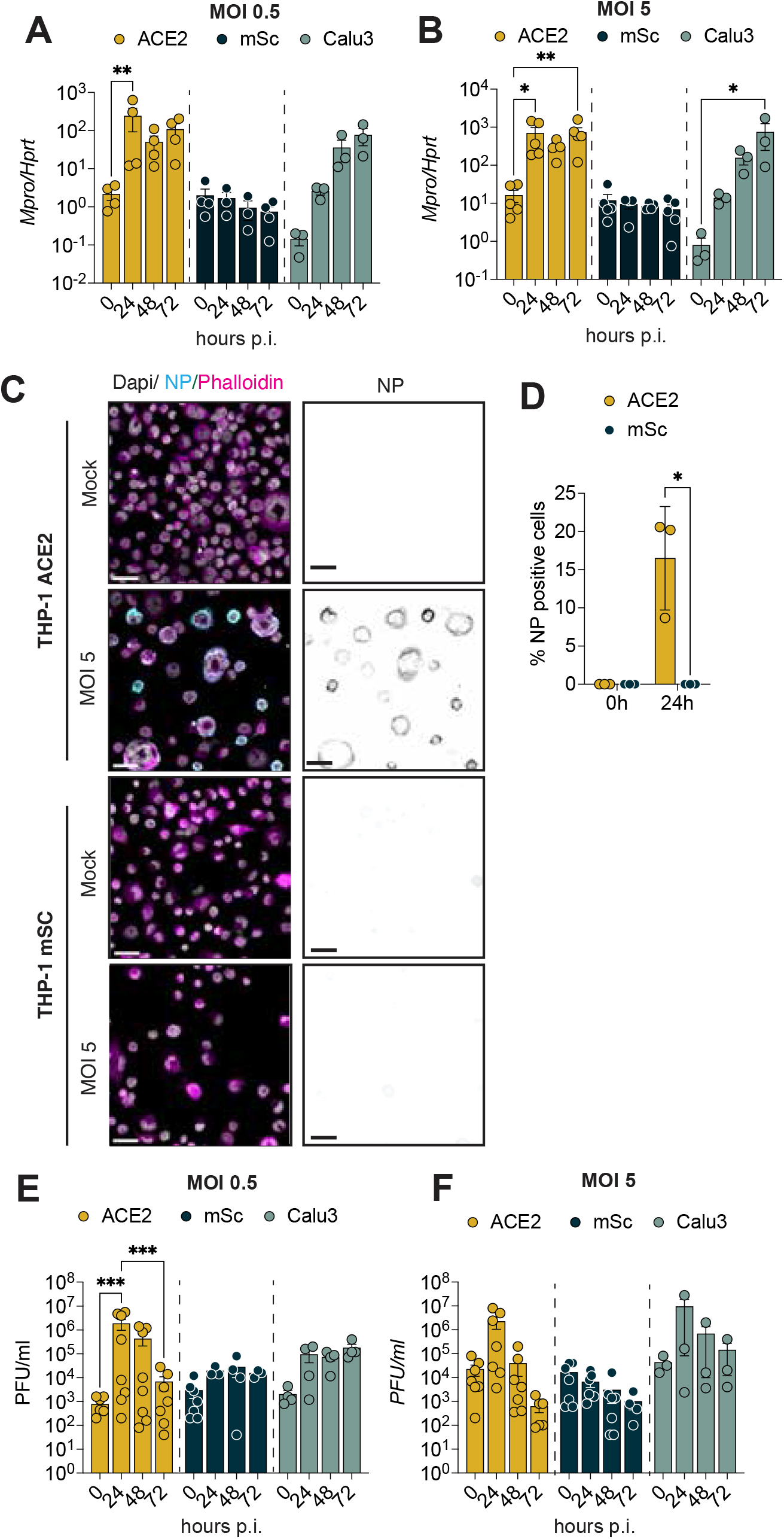
ACE2-positive macrophages support SARS-CoV-2 entry, early-stage replication and productive virion release. A-F: Cells were infected with SARS-CoV-2 at MOI 0.5 or MOI 5, as indicated. After one h, the virus inoculum was removed, cells were washed, and cells or supernatants were harvested at the indicated times (0 h = immediately after the virus inoculum was removed). Cellular viral mRNA was analyzed by qPCR (A-B). Viral N protein at 24 h was assessed by immunofluorescence staining, scale bar = 30 µm (C), and quantified relative to the total number of cells (D). Infectious virions released into cell supernatants were measured by plaque assay (E-F). Data show the mean + SEM of 3-8 independent experiments, with data points representing individual experiments. Asterisks indicate significance: p ≤ 0.05 (*), p ≤ 0.001 (**), p ≤ 0.0001 (***) (A-B, E-F: two-way ANOVA, Tukey’s multiple comparison test; D: unpaired t-test, Holm-Šídák test).

To determine whether the increase in cell-associated viral RNA levels in THP-1-ACE2 cells was coupled to the release of infectious virus, we performed plaque assays on cell-free supernatants between 0 h and 72 h p.i. At MOI 0.5, we observed a significant increase in infectious viral particles in the THP-1-ACE2 cell-free supernatants at 24 h p.i., compared with the levels of residual input virus (0 h p.i.), which is consistent with productive infection (**Fig. 3E**). We observed a similar increase in THP-1-ACE2 cells at 24 h p.i. using MOI 5 (**Fig. 3F**). Control epithelial cells (Calu-3: **Fig. 3E, 3F**, and A549-ACE2 cells, **Fig. S3**) supported productive virus release. As expected, there was no evidence of productive replication in control THP-1-mSc cells at either MOI (**Fig. 3E, 3F**), which is consistent with a lack of susceptibility to infection (**Fig. 3A, 3B**). This shows that ACE2-expressing macrophages support the initial stages of infection, new viral RNA synthesis, and assembly and release of newly synthesized, infectious viral particles.

### Macrophages can internalize SARS-CoV-2 independently of ACE2

Additional SARS-CoV-2 entry receptors beyond ACE2 are reported, including the C-type lectin receptors (CLRs) (*26*) and CD169(*33*). Since our assays to measure viral protein and RNA could not distinguish whether the virus is intracellular or extracellular (**Fig. 2C,2D**), we next determined whether SARS-CoV-2 can still bind and enter HMDM, despite the lack of ACE2 expression. We used transmission electron microscopy to determine the sub-cellular location of incoming virions, using a high MOI of 20. HMDM (**Fig. 4A, B**) and THP-1-ACE2 (**Fig. 4C**) displayed internalized, intact virions within the phagosomal system after one hour of infection. The THP-1-ACE2 cells also showed virions bound to the plasma membrane and possibly undergoing fusion (**Fig. 4D, E**). These virions were morphologically similar to the new structures released from Calu-3 at 72 h p.i. (**Fig. 4F**). Since THP-1-ACE2 cells endogenously express TMPRSS2 (**Fig. 1B**), this suggests that THP-1-ACE2 cells support viral fusion at the plasma membrane to deliver the viral genome and NP directly into the cytoplasm. Together, these results indicate that while HMDM take up SARS-CoV-2 into phagosomal compartments, low ACE2 expression will preclude SARS-CoV-2 S processing and virus-cell membrane fusion, necessary steps for this virus to enter the cytoplasm.

**Figure 4:**
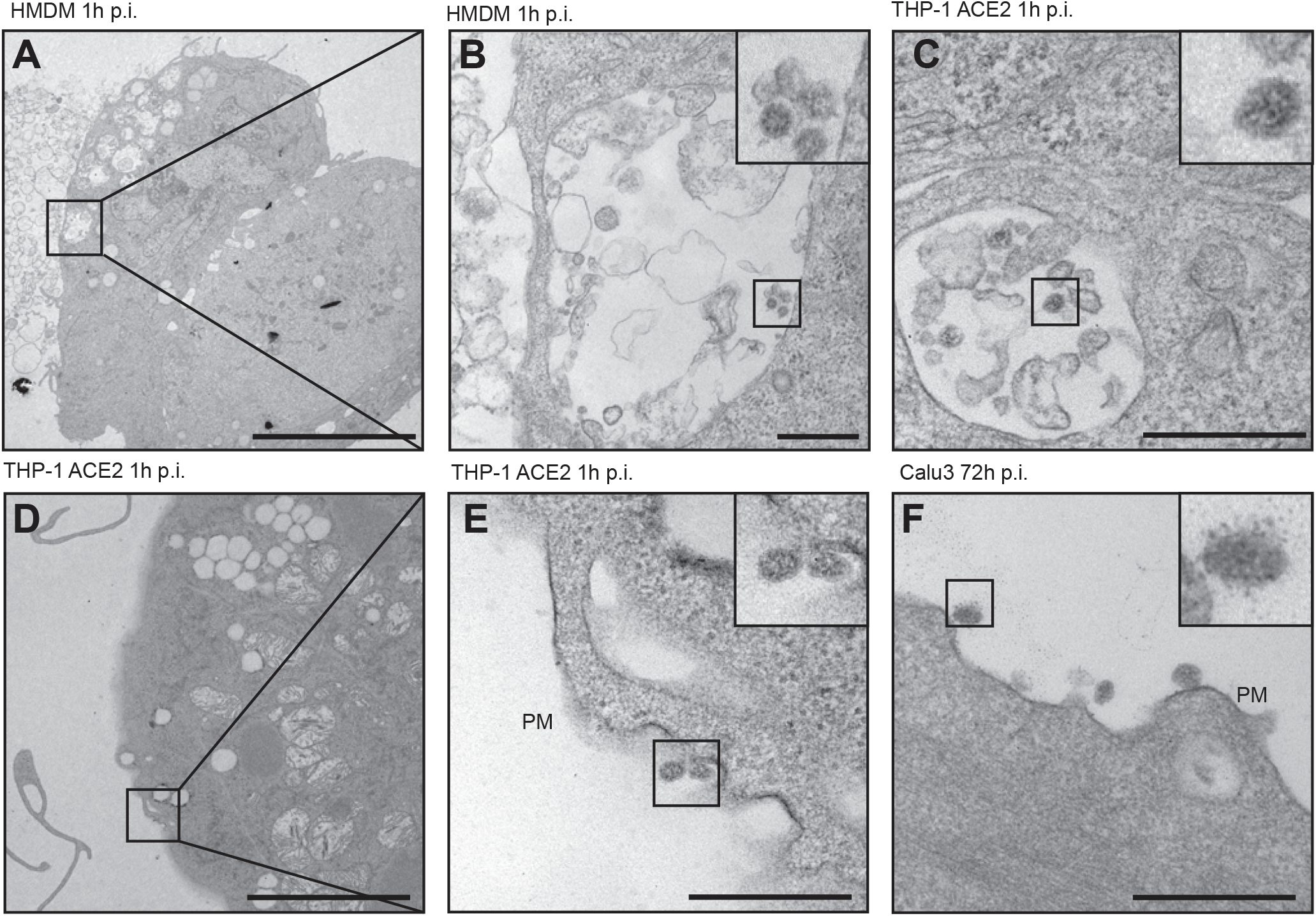
Macrophages can take up SARS-CoV-2 independently of ACE2. A-F: Transmission electron microscopy of indicated cells infected with SARS-CoV-2 (MOI 20) at indicated time points. For low magnification images (A, D) scale bar = 10 µm, and for all other images (B, C, E, F) scale bar = 500 nm.

### ACE2-negative macrophages do not release pro-inflammatory cytokines or anti-viral mediators upon SARS-CoV-2 exposure

Macrophages express a suite of pattern recognition receptors (PRRs) at strategic subcellular locations to detect microbial challenges. PRRs can sense viral RNA and proteins incorporated in incoming virions and RNA and proteins synthesized during viral replication (*34–36*). We detected viral particles in HMDM phagosomal compartments at one-hour p.i. (**Fig. 4B**), and so we next assessed whether ACE2-negative macrophages could sense incoming virions from these compartments. We challenged HMDM with high dose SARS-CoV-2 (MOI 5) for 24 h without removing the virus to allow maximal macrophage responses. SARS-CoV-2 did not trigger cytokine (C-X-C motif chemokine ligand 10: CXCL10; Interleukin-6: IL-6; or Tumor Necrosis Factor: TNF) release in vitro (**Fig. 5A**), despite reports of high circulating levels of these cytokines during human SARS-CoV-2 infection (*37, 38*). By contrast, HMDM showed a robust secretory response to synthetic viral mimetics such as the toll-like receptor (TLR) 7/8 ligand, R848 (TNF, IL-6), and the melanoma differentiation-associated protein 5 (MDA5) ligand, transfected poly I:C (pIC) (CXCL10) (**Fig. 5A**). mRNA analyses revealed that SARS-CoV-2 did not provoke HMDM expression of *IFNB1, IFNL1, CXCL10, IL6, TNF*, or *IL1B* at two or 24 h p.i., at either low (0.5) or high (5) MOI (**Fig. 5B)**. R848 and pIC stimulation robustly induced these genes, indicating that the HMDM were signaling competent (**Fig. 5B**). Thus, HMDMs do not respond to SARS-CoV-2 exposure in vitro, even at the high MOI of 5 or when virions are present in the phagosomal system.

**Figure 5:**
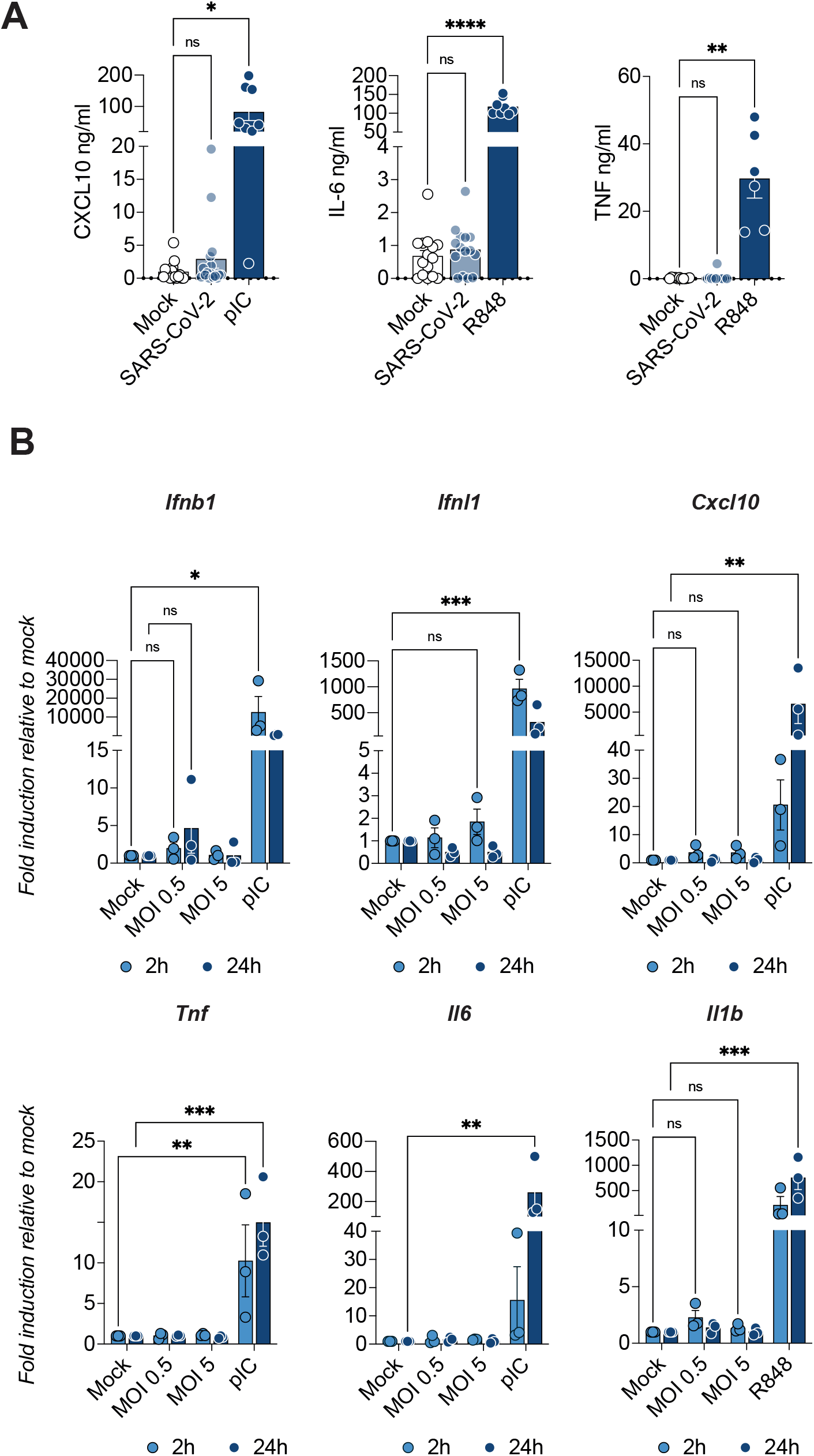
SARS-CoV-2 does not trigger inflammatory responses in ACE2-negative HMDM. A: HMDM were infected with SARS-CoV-2 (MOI 5) or stimulated with R848 or transfected pIC for 24 h. Cytokines in cell supernatants were analyzed by alphaLISA. Each data point represents an individual donor (n = 6-15). Graphs show mean + SEM, and significance is indicated by asterisks (one-way ANOVA, Dunnett’s multiple comparison test). B: HMDM were challenged with SARS-CoV-2 (MOI 0.5, 5) or transfected pI:C. Gene expression at 2 and 24 h was analyzed by qPCR. Graphs show mean + SEM, where each data point represents an individual donor (n = 3), and asterisks indicate significance: p ≤ 0.05 (*), p ≤ 0.001 (**), p ≤ 0.0001 (***) (two-way ANOVA, Dunnett’s multiple comparison test).

### ACE2-positive cells sense replicating SARS-CoV-2 to trigger pro-inflammatory and anti-viral responses

Since ectopic ACE2 expression in macrophages permits efficient SARS-CoV-2 entry and early-stage viral replication, we examined whether THP-1-ACE2 cells produce pro-inflammatory and anti-viral mediators upon SARS-CoV-2 challenge. THP-1-ACE2 cells strongly upregulated *IFNB1, IFNL1, CXCL10*, and *IL6* mRNA expression after infection with SARS-CoV-2 for 24 h (at MOI 5, **Fig. 6A**; or MOI 0.5, **fig. S4A**), correlating with increased SARS-CoV-2 viral RNA levels (**Fig. 3A-B**). In contrast, THP-1-mSc cells did not respond to SARS-CoV-2 challenge at either MOI (**Fig. 6A, fig. S4A)**, like HMDM (**Fig. 5A, 5B**). SARS-CoV-2 infection downregulated *IL1B* mRNA levels in THP-1-ACE2 cells but not in THP-1-mSc cells (**Fig. 6B, fig. S4A**), likely due to negative regulation of *IL1B* mRNA expression by interferons (*39*). Viral sensing pathways were operational in THP-1-ACE2 and THP-1-mSc cells, as both cell types responded equally to MDA5 stimulation via pIC transfection, including robustly downregulating *IL1B* mRNA expression (**fig. S4B**). In Calu-3 cells, anti-viral (*IFNB1, IFNL1, CXCL10*) and pro-inflammatory (*TNF, IL6*) gene induction peaked at 72 h p.i. (at MOI 5, **Fig. 6C**; MOI 0.5, **fig. S4C**), consistent with published observations (*7*) and the rise in SARS-CoV-2 viral RNA expression (**Fig. 3A, B**) and productive virus release (**Fig. 3E, F**). To determine whether THP-1-ACE2 cells responded to incoming versus replicating virus, we treated cells with the RdRp inhibitor remdesivir (*40*). Remdesivir ablated viral replication (**Fig. 6D**) and the ensuing induction of *IFNB1, IFNL1*, and *IL6* (**Fig. 6E**). Intriguingly, remdesivir did not modulate *CXCL10* expression during infection at MOI 5 (**Fig. 6E**). This suggests that incoming virions may also be sensed and trigger distinct inflammatory responses compared with replicating virions. Collectively, these data indicate that ACE2-positive macrophages are poised to sense entry and early-stage viral replication and respond by mounting potent anti-viral programs.

**Figure 6:**
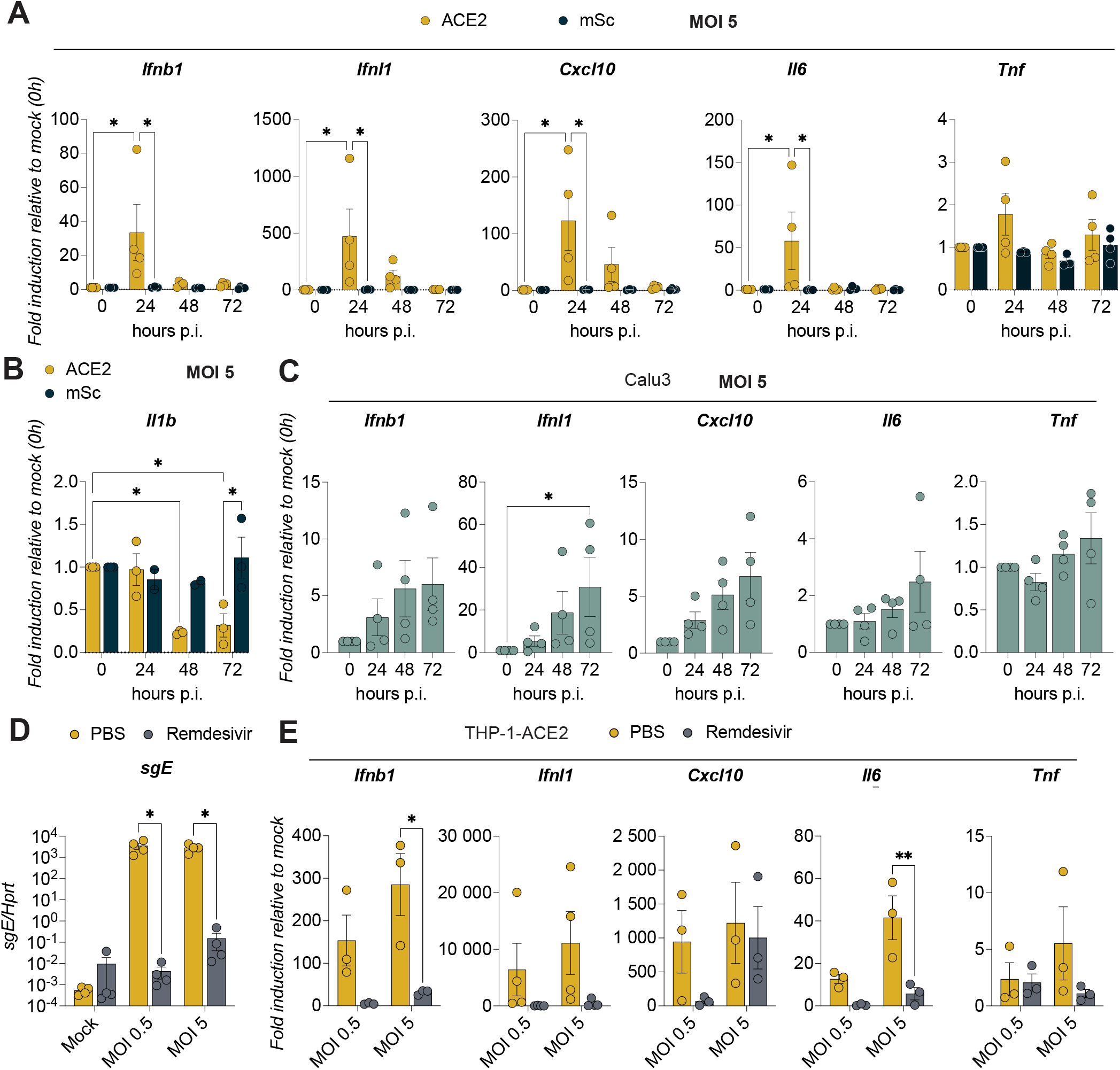
ACE2-positive macrophages sense entering or replicating SARS-CoV-2 to drive pro-inflammatory and anti-viral responses. A-C: THP-1-ACE-2 and THP-1-mSc cells (A,B) or Calu-3 cells (C) were infected with SARS-CoV-2 at MOI 5. Cells were harvested at the indicated times (0 h = immediately after the virus inoculum was removed), and gene expression was quantified by qPCR. Gene expression at each time point is presented relative to the mock control to show fold-gene induction. Data are mean + SEM of 4 independent experiments (indicated by individual data points), and significance is indicated by asterisks: p ≤ 0.05 (*), p ≤ 0.001 (**), p ≤ 0.0001 (***) (two-way ANOVA, Tukey’s multiple comparison test). (D-E). THP-1-ACE2 cells were infected with SARS-CoV-2 at MOI 5 in the presence of 10 µm remdesivir, and viral RNA (C) or inflammatory cytokines (D) were measured at 24 h p.i. Data show mean + SEM of 3 independent experiments (indicated by individual data points), and asterisks indicate significance: p ≤ 0.05 (*), p ≤ 0.001 (**), p ≤ 0.0001 (***) (two-way ANOVA, Sidak’s multiple comparison test).

### Blocking cytokine signaling in ACE2-positive cells prolongs productive virion release

Recent studies suggest that host cell death curbs the release of SARS-CoV-2 infectious virions (*41*). We next explored whether the death of THP-1-ACE2 cells was responsible for the decline in viral titers at 48h and 72h p.i. (**Fig. 3E, 3F**). Indeed, virus-challenged THP-1-ACE2 cells showed a modest but significant decrease (∼20%) in viability compared to THP-1-mSc cells at 72 h p.i. (**Fig. 7A**). Thus, cell death may limit virus release from THP-1-ACE2 challenged with high viral loads.

**Figure 7:**
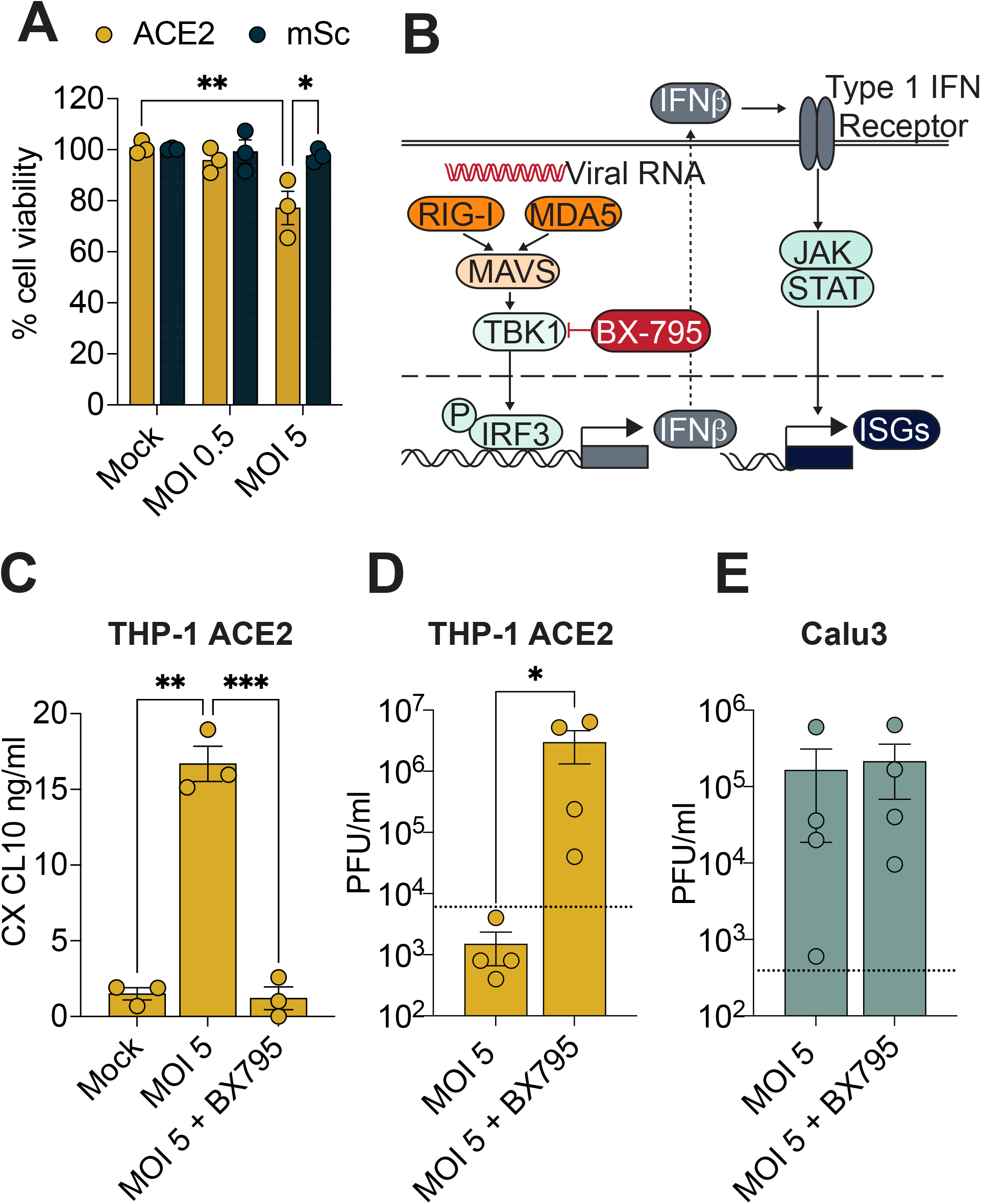
Blocking cytokine signaling in ACE2-positive macrophages prolongs infectious virion release. A: THP-1-ACE-2 and THP-1-mSc cells were infected with SARS-CoV-2 (MOI 5). Cell death was analyzed by ATPlite assay at 72 h. Data are presented as cell viability relative to mock infection and are mean + SEM of 3 independent experiments. Asterisks indicate significance: p ≤ 0.05 (*), p ≤ 0.001 (**), p ≤ 0.0001 (***) (two-way ANOVA, Tukey’s multiple comparison test). B: Schematic of TBK1 (BX-795) inhibition. C-E: THP-1-ACE-2 or Calu-3 cells were stimulated with SARS-CoV-2 at MOI 5. After one h, the viral inoculum was removed, and BX-795 was added. Supernatants were harvested at 72 h, CXCL10 was analyzed by ELISA (C) and viral titers were analyzed by plaque assay (D, E). Data show the mean + SEM of at least three independent experiments, with each data point representing a different experiment. Significance is indicated by asterisks: p ≤ 0.05 (*), p ≤ 0.001 (**), p ≤ 0.0001 (***) (C: one-way ANOVA, Tukey’s multiple comparison test; D, E: ratio-paired t-test).

We also noted that the rapid and robust induction of type I and III interferons in THP-1-ACE2 cells peaked at 24 h p.i., much earlier than in Calu-3 cells (**Fig. 6A, 6B**). Interferons provide autocrine and paracrine signals to activate an anti-viral state through the induction of interferon-stimulated genes, which act as restriction factors to limit viral infection (*42*). Indeed, type I IFN aborts influenza A virus infection in murine macrophages (*16*). We thus hypothesized that interferon signaling in THP-1-ACE2 cells induces host factors that impede viral replication, limiting the release of productive virions. In contrast, the delayed induction of *IFNB1* and *IFNL1* in Calu-3 cells, potentially through viral antagonism, may contribute to ongoing viral replication and release at 48 and 72 h p.i. (**Fig. 3E, 3F**). We tested this hypothesis using a TBK1 inhibitor (BX-795) to block virus-induced interferon induction and signaling (*43*) in THP-1-ACE2 cells (**Fig. 7B**). As expected, BX-795 suppressed pIC-induced expression of *IFNB1, IFNL1*, and the interferon-stimulated gene *CXCL10* (**fig. S5**). We next challenged THP-1-ACE2 and Calu-3 cells with SARS-CoV-2 for 72 h in the presence and absence of BX-795. BX-795 suppressed SARS-CoV-2-induced CXCL10 production from THP-1-ACE2 cells (**Fig. 7C**), significantly increasing viral release at 72h p.i. (**Fig. 7D**). BX-795 did not boost the release of productive virions from Calu-3 cells (**Fig. 7E**), consistent with published observations (*7*). These findings indicate that even though SARS-CoV-2 can enter ACE2-positive macrophages, rapid anti-viral signaling limits viral replication and release.

## DISCUSSION

A key question in SARS-CoV-2 pathogenesis is which type of host cells efficiently sense SARS-CoV-2 to trigger inflammatory cytokine and anti-viral mediator release. Since macrophages are prime candidates for sensing and responding to SARS-CoV-2 (*11*), a second unresolved question is whether SARS-CoV-2 can infect and productively replicate in human macrophages. This study identifies ACE2 as a critical determinant of macrophage susceptibility to infection and sensing of infectious SARS-CoV-2.

We demonstrate here that HMDM do not express ACE2 and are thus suitable in vitro models for ACE2-negative macrophages. While HMDMs phagocytose virus, they do not permit early-stage viral replication or protein synthesis and do not sense phagocytosed virus to trigger inflammatory responses. Our observation that SARS-CoV-2 does not replicate in HMDM or BAL is consistent with multiple reports of abortive SARS-CoV-2 infection of macrophages in vitro (*12, 13, 23, 24*). While these studies demonstrate that macrophage viral challenge results in decreasing viral RNA (*12, 13, 23*) and viral protein (*24*) levels, and macrophages fail to produce new infectious virus (*12, 23*), these reports nevertheless observe virus entry into macrophages via quantification of SARS-CoV-2 nucleoprotein-positive cells. This contrasts with studies reporting no viral entry into HMDM (*7, 25*). We observe virions within phagosomes (**Fig. 4B**), indicating that SARS-CoV-2 virions can enter the macrophage phagolysosomal system. Despite this, we propose that the virus does not enter the HMDM cytoplasm, likely because, without ACE2, the Spike protein does not undergo the necessary conformational changes for membrane fusion.

ACE2 expression analysis across cell types in healthy and COVID-19 lungs shows that a small proportion of macrophages express ACE2, corroborating new reports that both ACE2-positive and -negative macrophages are likely to be present at sites of infection in vivo (*20, 27, 29*). Age, sex, and co-morbidities may influence cellular ACE2 expression (*44*), as well as environmental cues such as hypoxia within tissues (*45, 46*). How macrophage ACE2 expression is regulated has yet to be elucidated. Uncovering the macrophage-specific cues and pathways regulating ACE2 expression and function will yield further insight into how ACE2-positive macrophages can be detected and targeted.

In line with our findings herein, other non-epithelial cells that do not usually express ACE2, such as endothelial cells, are rendered susceptible and permissive to productive SARS-CoV-2 infection by ectopic ACE2 expression (*32*). In THP-1 cells, ectopic ACE2 expression facilitated SARS-CoV-2 entry and early-stage replication, but in response, these cells initiated an interferon program that limits new virion release after 24h p.i. (**Fig. 3E, 3F, 6A**). This limit could reflect the death of infected macrophages, as observed for antibody-dependent monocyte infection(*47*), or restriction by macrophage anti-viral programs. While we observe some virus-induced cell death in ACE2-positive macrophages (**Fig. 7A**), our TBK1 inhibitor experiments (**Fig.s 7C-D**) indicate that a potent macrophage anti-viral response interrupts the viral replication cycle. Future studies should elucidate the precise anti-viral programs underpinning macrophage restriction of SARS-CoV-2. Nevertheless, our findings suggest that ACE2-positive macrophages induce a sufficiently robust anti-viral response to prevent them from acting as ‘Trojan horses’ to disseminate SARS-CoV-2 to extra-respiratory tissues. If such anti-viral signaling programs are compromised, such as in patients with inborn errors of immunity or circulating IFN autoantibodies(*48, 49*), then ACE2-positive macrophages may substantially contribute to increased viral loads and viral dissemination.

The proportion of macrophages positive for viral RNA in COVID-19 lungs is much higher than that of ACE2-positive macrophages (*20, 50, 51*). Our data suggest that this is not due to ACE2-independent SARS-CoV-2 entry into and replication within macrophages, in contrast to a recent study that reported CD169-dependent, ACE2-independent macrophage infection(*33*). Instead, we expect macrophages positive for viral RNA may be ACE2-negative macrophages with phagocytosed virions (**Fig. 4B**), extracellular viral RNA, or virus-infected cells (*52*). Alternatively, ACE-negative macrophages might be infected via an antibody-dependent uptake route, as recently shown for SARS-CoV-2-infected human monocytes (*47*). Our data show that the small proportion of macrophages positive for ACE2 can be directly infected and trigger a distinct inflammatory response to ACE2-negative macrophages encountering SARS-CoV-2 virions. Importantly, for ACE2-positive macrophages, our data suggest that the viral RNA present in macrophages in vivo also reflects limited productive infection. A direct comparison of macrophage responses to these distinct routes of viral uptake would elucidate whether targeting one subset could maintain protective anti-viral responses while potentially limiting pathology.

The failure of primary HMDM to respond to SARS-CoV-2 exposure (**Fig. 5A, 5B**) contrasts with previous reports that SARS-CoV-2 induces the expression of cytokine mRNAs (including *IL6, CXCL10*, and *TNF*) in HMDM (*12, 13, 26, 53*). Typically, a virus must enter and replicate within a cell to activate cytosolic PRRs (so-called ‘cell intrinsic’ sensing), concomitant with a high threat to that cell. Our data indicate that ACE2-negative macrophages do not respond directly to SARS-CoV-2 exposure with cytokine production (**Fig. 5A, 5B**) because lack of virus entry precludes ‘cell intrinsic’ sensing. Our results thus agree with published observations that SARS-CoV-2 does not trigger cytokine production from HMDM (*25*) and that SARS-CoV-2 also fails to trigger the interferon system in human alveolar macrophages (*24*). In contrast, ACE2-positive macrophages detected actively replicating SARS-CoV-2 to trigger potent *IFNB1* and *IFNL1* induction (**Fig. 6E**). We hypothesize that the cytosolic viral sensors retinoic acid-inducible gene-1 (RIG-I) and MDA5, which detect replicating SARS-CoV-2 RNA in epithelial cells (*7, 54, 55*), drive macrophage anti-viral responses. Macrophage-derived interferon critically orchestrates the anti-viral immune response to limit disease in other respiratory viral infections (e.g., Newcastle Disease Virus, Respiratory Syncytial Virus) (*17, 18*). We speculate that interferon produced by ACE2-positive macrophages via cell-intrinsic sensing may also contribute to early control of SARS-CoV-2 infection in the lung. Alternatively, this same macrophage-derived interferon could contribute to disease pathology (*56, 57*).

In addition to cytosolic PRRs that detect serious threats, macrophages are also equipped with PRRs at other subcellular locations (e.g., cell surface, endosomes) to detect moderate threats. This ‘cell extrinsic’ sensing detects neighboring infected cells, viral pathogen-associated molecular patterns (PAMPs), or host danger-associated molecular patterns (DAMPs) in the extracellular milieu (*58*). We and others (*24, 25*) find that HMDM do not sense SARS-CoV-2, either at the cell surface or in endosomes, to trigger pro-inflammatory responses (**Fig. 5**), which contradicts reports that SARS-CoV-2 selectively induces a pro-inflammatory response (e.g., TNF, IL-6, CXCL10) from HMDM (*12, 13, 23*). Differences in viral stock preparation or quantification could underpin these divergent observations. SARS-CoV-2 is usually cultured in cell lines (e.g., VERO E6 or Calu-3 cells (*12*)) that inherently respond to infection by producing cytokines or undergoing cell death. Indeed, virus is often harvested at a time point when a visible cytopathic effect emerges. Individual viral preparations quantified for intact virions by TCID_50_ or plaque assays may also contain free viral RNA, viral proteins, non-infectious virions, and parental cell line-derived DAMPs and cytokines (*59*). Thus, published macrophage responses to viral preparations could represent an indirect response to epithelial DAMPs or free viral PAMPs. Nevertheless, our viral stocks did not trigger cytokine release from HMDM or THP-1 cells in the absence of ACE2, indicating that ACE-dependent entry and early-stage replication are necessary for macrophage sensing (**Fig. 5A, 5B, Fig. 6A**). We anticipate this observation will also apply to ACE2-dependent SARS-CoV-2 variants, including Omicron (BA.4 and BA.5 lineages) and, indeed, any beta-coronaviruses that utilize ACE2. Consistent with this, SARS-CoV-1, which also requires ACE2 for entry, does not trigger macrophage cytokine responses (*60*).

This study reveals that human macrophages do not sense SARS-CoV-2 unless they express ACE2 and support new viral synthesis. Our findings give new insight into SARS-CoV-2 cell tropism and the influence therein of macrophage innate immune pathways. Such studies of innate immune cell-intrinsic and -extrinsic SARS-CoV-2 recognition and response will help reveal novel therapeutic targets for new drugs that dampen pathogenic pro-inflammatory signaling in virulent viral infections without impeding host anti-viral defense.

## MATERIALS AND METHODS

### Re-analysis of human lung scRNA-seq

We reanalyzed three lung scRNA-seq datasets representing 1) samples from control or COVID-19-infected individuals (*27*), 2) samples from COVID-19-infected individuals (*20*), and 3) samples from control individuals (*28*).

To compare matched cell types across datasets, we used Seurat v 4.1.1(*61*) integration and label transfer analysis to impose the ‘fine level’ cell identities defined by Melms et al. onto the remaining datasets. We first applied the Seurat CCA integration pipeline to integrate the individuals from Melms et al. before label transfer analysis. Functions *NormalizeData* and *FindVariableFeatures* were run on each individual, and *SelectIntegrationFeatures* was run with *nfeatures* set to 1000. *FindIntegrationAnchors* and *IntegrateData*, followed by *ScaleData*, and *RunPCA*, were all run with default parameters. We then transferred the cell type identities from Melms et al. to the remaining datasets using the *FindTransferAnchors* and *TransferData* functions with dims = 1:50.

### Reagents and inhibitors

Low molecular weight Poly I:C (Invivogen, tlrl-picw) was transfected into cells with lipofectamine LTX (Thermofisther, A12621) at 0.5 µg pIC per well, while R848 Resiquimod (Miltenyi Biotech, 130-109-376) was used at a final concentration of 250 ng/ml. The TBK1 inhibitor BX-795 (Sigma-Aldrich, SML0694) was used at a final concentration of 5 µM. Remdesivir (Gilead) at a final concentration of 10 µM.

### Cells

Peripheral blood mononuclear cells were isolated from buffy coats by density centrifugation using Ficoll-Paque Plus (GE Healthcare). CD14+ monocytes were subsequently isolated using magnetic-activated cell sorting (Miltenyi Biotech) according to the manufacturer’s instructions. Human macrophages were differentiated from human CD14+ monocytes as previously described (*62*) and then used for experiments on day 7 of differentiation. Human monocyte-derived macrophages (HMDMs) were cultured in media consisting of RPMI 1640 medium (Life Technologies) supplemented with 10% heat-inactivated fetal bovine serum (FBS), 2 mM GlutaMAX (Life Technologies), 50 U/ml penicillin-streptomycin (Life Technologies) and 150 ng/ml recombinant human macrophage colony-stimulating factor (CSF-1, endotoxin-free, expressed and purified by the University of Queensland Protein Expression Facility). HMDM were seeded 16 h prior to experiments at 500 000 cells per well in 12-well plates or 200 000 cells per well in 24-well plates. Studies using primary human cells were approved by the University of Queensland Human Medical Research Ethics Committee. THP-1 cells (TIB-202; ATCC) were maintained in RPMI 1640 medium supplemented with 10% heat-inactivated fetal bovine serum (FBS), 2 mM GlutaMAX (Life Technologies) and 50 U/ml penicillin-streptomycin. For experiments, THP-1 cells were seeded at 500 000 cells per well of a 12-well plate and differentiated for 48 h with 30 ng/ml phorbol-myristate-acetate (PMA). Calu-3 cells purchased from ATCC (HTB-55) were maintained in Minimal Essential Media (Invitrogen) containing 10% heat-inactivated fetal bovine serum (Cytiva), 50 U/ml penicillin, and streptomycin (Life Technologies Australia), and then seeded at 300 000 cells per well in 12-well plates 48 h prior to experiments. A549 cells (provided by Dr. Chris Macmillan, SCMB, UQ) were maintained in Dulbecco’s Modified Essential Medium (DMEM; Invitrogen) supplemented with 10% heat-inactivated FBS and 50 U/ml penicillin-streptomycin.

Patient bronchoalveolar lavage (BAL) was obtained at the time of diagnostic bronchoscopy as previously described (*63*). Briefly, the bronchoscope was wedged into a non-dependent sub-segmental bronchus (*64*) of a radiologically normal segment of the lung, and 20 ml of normal saline was instilled, retrieved, and then discarded to clear the bronchoscope of bronchial secretions. A further 80-100 ml was instilled in 20 ml aliquots and retrieved via hand aspiration of the syringe. Studies using primary human cells were approved by the Royal Melbourne Hospital and the University of Melbourne Human Research Ethics Committees. BAL was filtered, and cells were washed and seeded at 1×10^6^ cells per ml overnight in 48-well tissue culture plates in RPMI 1640 medium supplemented with 10% heat-inactivated fetal bovine serum (FBS), two mM GlutaMAX (Life Technologies) and 50 U/ml penicillin-streptomycin (Life Technologies). Non-adherent cells were removed via media change four hours post-seeding, resulting in a >90% macrophage population, as described previously (*65*).

### Lentiviral transduction

A lentiviral construct containing human ACE2 (Addgene 155295) or mScarlet (Addgene 85044) was cloned into pLV-CMV-MCS-IRES-Puro-Sin (*66*) and packaged into lentivirus in HEK-293T cells using third-generation lentiviral packaging plasmids (*67*). Lentivirus-containing supernatant was harvested on days 2 and 3 after transfection and then concentrated using a Lenti-X concentrator (Clontech, 631232). HEK-293T cells were transfected with the expression vectors according to the manufacturer’s protocol with PEI 2500 (BioScientific), then transduced target THP-1 cells were selected with puromycin (1 µg/mL) after 24 h and used for assays after 72 h.

### Viruses and cell infections

SARS-CoV-2 isolate hCoV-19/Australia/QLD02/2020 was provided by Queensland Health Forensic & Scientific Services, Queensland Department of Health. Virus was grown on Vero E6 TMPRSS2 cells for 48 h in DMEM with 2% FBS, and cell debris was cleared by centrifugation at 500G for 5 minutes at room temperature. Virus was titered as described previously by plaque assay (*32*). Sanger sequencing was used to confirm that no mutations occurred in the spike gene relative to the original clinical isolate. Cells (HMDM, THP-1, Calu-3) in 12-well plates (500 000 cells/well) were challenged for 1 h at 37°C and 5% CO_2_ with 2.5 × 10^6^ plaque-forming units (PFUs) for MOI 5, or 2.5 × 10^5^ PFUs for MOI 0.5. For Calu-3 cells, virus was added to cells for a total volume of 500 µL of RPMI 1640 with 2% FBS (HMDM and THP-1) or MEM with 2% FBS (Calu-3) per well. The viral inoculum was removed, and the medium was replaced with DMEM (Invitrogen) or MEM (Invitrogen) containing 2% FBS. Alternatively, virus was not removed from the cells. All studies with SARS-CoV-2 were performed under physical containment 3 (PC3) conditions and were approved by The University of Queensland Biosafety Committee (IBC/374B/ SCMB/2020, IBC/518B/IMB/SCMB/2022) and the University of Melbourne Institutional Biosafety Committee in consultation with the Doherty Institute High Containment Facility Management Group.

For studies involving SARS-CoV-2 infection of BAL macrophages, the SARS-CoV-2 isolate hCoV-19/Australia/VIC01/2020 (kindly provided by the Victorian Infectious Diseases Reference Laboratory) was grown in Vero cells for 72 h in serum-free MEM with 1 µg/ml TPCK trypsin. The median tissue culture infectious dose (TCID_50_) was calculated using the Reed-Muench method. BAL macrophages (approx. 2.5 × 10^5^ cells) were seeded overnight in 48-well plates. Macrophages were infected with SARS-CoV-2 (MOI 1) in serum-free media (RPMI supplemented as described above) for 1 h. The inoculum was removed, and cells were washed before the media was replaced with 400 µl serum-free media for a further 2 to 48 h. VERO control cells were infected at an MOI of 0.5, and maintenance media included 1 µg/ml of TPCK trypsin. RNA from cell-free supernatant was collected and extracted using a QIAamp Viral RNA Mini Kit (Qiagen), while RNA from cell monolayers was extracted using the Rneasy Plus Mini Kit (Qiagen).

### Cell Death

THP-1 were seeded at 50 000 cells per well of a black-walled, clear-bottomed 96-well plate (CLS3916, Corning). After 48 h of differentiation with PMA, cells were inoculated with virus at MOI 5 and incubated for one h before the media was replaced with 100 µl of fresh RPMI with 2% FCS. After 72 h, ∼60 µl of media was removed per well, and 30 µl of ATPlite substrate solution (Perkin Elmer) was added to the remaining 30 µl. After 10 minutes of incubation at room temperature, plates were sealed with Optical Adhesive Film (Thermofisher), and luminescence was read on a Victor Nivo Plate Reader (Perkin Elmer).

### RNA extraction and qPCR analysis

Cells were lysed in Buffer RLT plus β-mercaptoethanol, and the RNA was directly processed using a Rneasy Mini Kit (Qiagen) with on-column DNase digestion according to the manufacturer’s instructions. RNA concentration was measured using a NanoDrop spectrophotometer, with equal starting concentrations of RNA for each sample used for reverse transcription. Reverse transcription was performed using Superscript III Reverse Transcriptase (ThermoFisher) with random hexamer priming. Quantitative PCR was performed using SYBR green reagent (Applied Biosystems) on a QuantStudio 7 Flex Real-Time PCR System (ThermoFisher) in 384-well plates (Applied Biosystems), and relative gene expression was determined using the change-in-threshold (2^-ΔΔCT^) method, using Hypoxanthine Phosphoribosyltransferase 1 (*HPRT*) as an endogenous control. Alternatively, gene expression was determined relative to a standard curve generated from plasmids containing SARS-CoV-2 Main Protease (Mpro). Primers are as follows: *HPRT* F: TCAGGCAGTATAATCCAAAGATGGT R: AGTCTGGCTTATATCCAACACTTCG; *ACE2* F: TCACGATTGTTGGGACTCTGC, R: TCGCTTCATCTCCCACCACT; *IL6* F: CTCAGCCCTGAGAAAGGAGACAT, R: TCAGCCATCTTTGGAAGGTTCA; *TNF* F: TGCCTGCTGCACTTTGGAGTGA, R: AGATGATCTGACTGCCTGGGCCAG; *IL1B* F: GAAGCTGATGGCCCTAAACA, R: AAGCCCTTGCTGTAGTGGTG, *IFNB1* F: CAGTCCTGGAAGAAAAACTGGAGA, R: TTGGCCTTCAGGTAATGCAGAA; *CXCL10* F: TGAAAGCAGTTAGCAAGGAAAGGT, R: AGCCTCTGTGTGGTCCATCC; *IFNL1* F: CGCCTTGGAAGAGTCACTCA, R: GAAGCCTCAGGTCCCAATTC; SARS-CoV-2 *Mpro* F: GAGACAGGTGGTTTCTCAATCG, R: ACGGCAATTCCAGTTTGAGC; SARS-CoV-2 *E* gene F: ACAGGTACGTTAATAGTTAATAGCGT, R: ATATTGCAGCAGTACGCACACA. *Subgenomic E* gene: F: AATATTGCAGCAGTACGCACACA, R: CGATCTCTTGTAGATCTGTTCTC

Copy number of the SARS-CoV-2 E gene was measured in BAL or VERO cellular RNA (equal starting concentration of RNA) and in RNA from cell culture media (equal starting volume). SensiFAST™ Probe Lo-ROX One-Step Kit with E-specific probe (FAM-ACACTAGCCATCCTTACTGCGCTTCG-QQA) was used to detect the E gene, and a plasmid-based standard curve was used to quantify the number of E copies. The following primers were used to amplify the Mpro sequence from cDNA with the Phusion polymerase kit (New England BioLabs). F:AATAAGGTACCAGTGGTTTTAGAAAAATGG, R:TTATTGCGGCCGCTCATTGGAAAGTAACACC.

The Mpro expression vector was generated by cloning the Mpro PCR product into a modified pEF6 plasmid, with an HA-tag N-terminal of the multiple cloning site, by standard restriction digest cloning techniques. The vector and correct insertion of Mpro were verified by Sanger sequencing.

### Cytokine analysis

Cytokine titers were determined using an AlphaLISA Immunoassay kit (Perkin Elmer) according to the manufacturer’s instructions and analyzed on a Victor Nivo Plate Reader (Perkin Elmer).

### Immunofluorescence

Cells were cultured on 12 mm glass coverslips and washed with PBS_+/+_ (PBS supplemented with 1mM CaCl2 and 0.5mM MgCl2, fixed for 30 min in 4% paraformaldehyde (Sigma Aldrich), and blocked and permeabilized for 30 min with 3% BSA, 0.3% Triton X-100 (Sigma Aldrich) at room temperature. Primary antibodies were incubated in 1.5% BSA for 1.5 h at room temperature, and secondary antibodies conjugated to Alexa fluorophores (Invitrogen) were incubated in 1.5% BSA for 60 min in the dark at room temperature. Cells were stained with Phalloidin (PHDN1-A; Jomar Life Research) and NP primary nanobody. NP primary nanobody was kindly supplied by Dr. Ariel Isaacs, Watterson Laboratory, School of Chemistry and Molecular Biology, UQ, QLD. After each incubation, cells were washed 3x with PBS_+/+_ and mounted with ProlongGold + DAPI solution (Cell Signaling Technologies) onto glass slides. Z-stack image acquisition was performed on a confocal laser scanning microscope (Zeiss LSM880) using a 40x NA 1.3 water immersion objective or 20x NA 0.8 objective. Raw images were processed and analyzed using Fiji (v1.53q). Images were batch processed using a custom-made FIJI script (*68*). The image processing included background subtraction and median filtering; then, the threshold was determined using the MinError filter. The holes were filled in the binary image, and Analyze Particles was used to detect and count nuclei and infected cells. Figures were prepared using Adobe Illustrator. For quantification analysis, five random regions of interest (ROIs) per group were acquired.

### Electron Microscopy

Electron microscopy samples were processed using a method adapted from a previous study (*69*). Briefly, cells were fixed in 2.5% glutaraldehyde (Electron Microscopy Services) in PBS for 24 h, then post-fixed in 1% osmium (ProSciTech) for one h and contrasted with 1% aqueous uranyl acetate (Electron Microscopy Services) for one h. Samples were then serially dehydrated in increasing percentages of ethanol before serial infiltration with LX-112 resin (Ladd Research) in a Biowave microwave (Pelco). Ultrathin sections were obtained using an ultramicrotome (UC6:Leica) and further contrasted using Reynold lead post-stain. Micrographs were acquired using a JEOL 1011 transmission electron microscope at 80 kV, with a Morada CCD camera (Olympus) utilizing iTEM software.

### Immunoblotting

For total cell lysates, cells were washed once with PBS and lysed with RIPA buffer (50 mM Tris, 150 mM NaCl, one mM EDTA, 1% Triton X-100, 0.1% SDS, 1% sodium deoxycholate, protease inhibitor, pH 8.0). A Pierce BCA protein assay kit (Thermo Scientific) was used to equalize protein amounts, wherein SDS sample buffer containing 100 mM DTT (Astral Scientific) was added, and samples were boiled at 95°C for 10 min to denature proteins. Proteins were then separated on 4-15% mini protean TGX precast gels (Bio-Rad) in running buffer (200 mM glycine, 25 mM Tris, 0.1% SDS (pH8.6)), transferred to PVDF or nitrocellulose membrane (Bio-Rad 1620112) in blot buffer (48 nM Tris, 39 nM glycine, 0.04% SDS, 20% MeOH) and subsequently blocked with 5% (w/v) milk powder or BSA in Tris-buffered saline with Tween-20 (TBST) for 30 min. Primary antibodies were incubated overnight at 4 deg C, followed by secondary antibodies linked to horseradish peroxidase (HRP) (Cell Signalling) or AlexaFluor647 (Invitrogen), and after each step, immunoblots were washed 3x with TBST. HRP signals were visualized by enhanced chemiluminescence (ECL) (Bio-Rad) and imaged with a Vilber Fusion Imaging system (Vilber). The fluorescence signal was detected using the AI600 imager (Amersham). The following antibodies were used: SARS-CoV-2 Nucleoprotein (Sino Biological), Tubulin (9F3; Cell Signalling Technology), ACE2 (AF933; RnD Systems), TMPRSS2 (Abcam) and Actin (8H10D10; Cell Signalling Technology), ACE2 (MA532307, Invitrogen), Calnexin (ab22595, Abcam).

### Statistical Analysis

Statistics were calculated using GraphPad Prism using tests as indicated in figure legends.

## Supplementary Materials

Figure S1: Re-analysis of macrophage ACE2 expression in vivo

Figure S2: SARS-CoV-2 does not replicate in HMDM

Figure S3: SARS-CoV-2 replicates in ACE2-A549 cells

Figure S4: Macrophage ACE2 expression potentiates macrophage inflammatory responses to SARS-CoV-2 at MOI 0.5

Figure S5: BX-795 blocks anti-viral cytokine induction in THP1-ACE2 cells.

## ACKNOWLEDGEMENTS

We gratefully acknowledge Dr. Lilian Schimmel for producing the ACE2 and mScarlet expressing lentiviruses, Dr. Ariel Isaacs for producing the anti-N nanobody, and Dr. Fiona Wylie for critical reading of this manuscript. This work was supported by National Health and Medical Research Council (NHMRC) of Australia funding, including fellowships (1124612 to LL; 1177174 to K. Subbarao; 1156489 to RGP; 2009075 to K. Schroder) and grants (2010757 to LL and EG; 2013574 to CMN, 1140064 and 1150083 to RGP; 2007979 to KRS; 1184532 to SL; 2009677 to K. Schroder) and the Doherty Institute COVID-19 Agility Fund. The Melbourne WHO Collaborating Centre for Reference and Research on Influenza is supported by the Australian Government Department of Health.

## AUTHOR CONTRIBUTIONS

Conception: LIL and SLL

Data acquisition and analysis: LIL, KYC, KE,XW, TE, CJS, JR, RP, HM, BH, TY, CLH, SE, SF, FLM, DPS, CMN, AKL, EG, RGP, SLL

Funding acquisition: LIL, SLL, K.Schroder

Manuscript original draft: LIL, K.Schroder

Manuscript editing: LIL, K.Subbarao, KRS, SLL, K.Schroder

## DATA AND MATERIALS AVAILABILITY

All data are available in the main text or the supplementary materials.

