## Supplementary materials for "Macrophages only sense infectious SARS-CoV-2 when they express sufficient ACE2 to permit viral entry, where rapid cytokine responses then limit viral replication"

**Supplementary Figure 1**

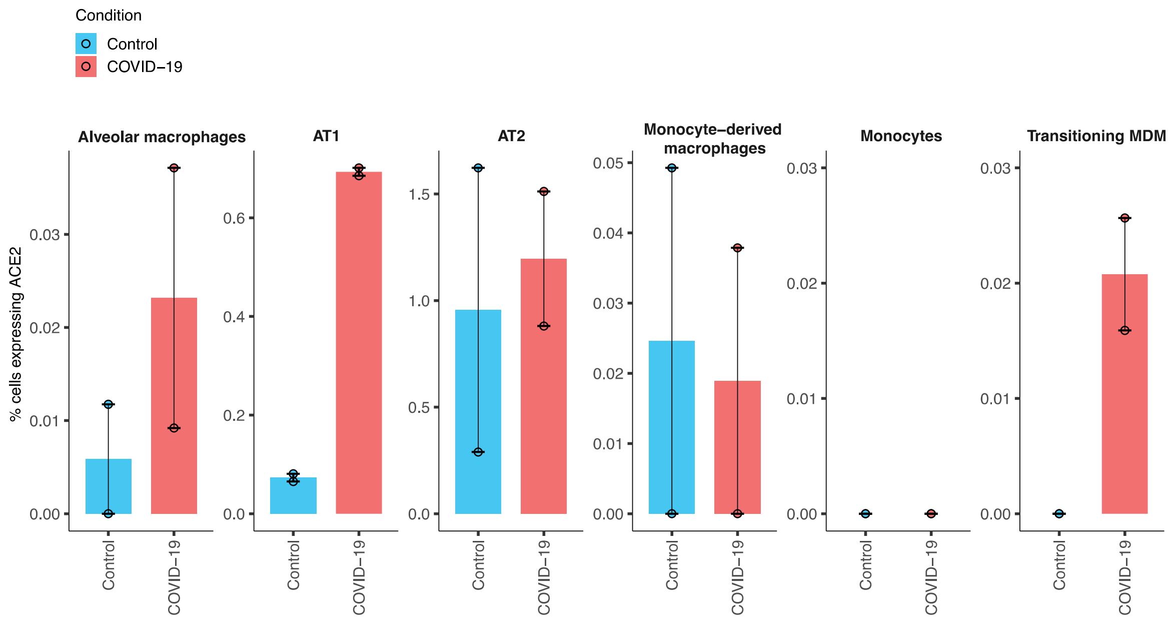


Figure S1: Re-analysis of macrophage ACE2 expression in vivo

Data shows the average percentage of cells expressing *ACE2* mRNA across indicated cell types. Data is from multiple lung scRNA-seq datasets representing two uninfected (control) and two COVID-19-infected cohorts. Datapoints represent the average of a cohort, with bars and error bars showing the overall average and SEM, respectively.

**Supp Figure 2**

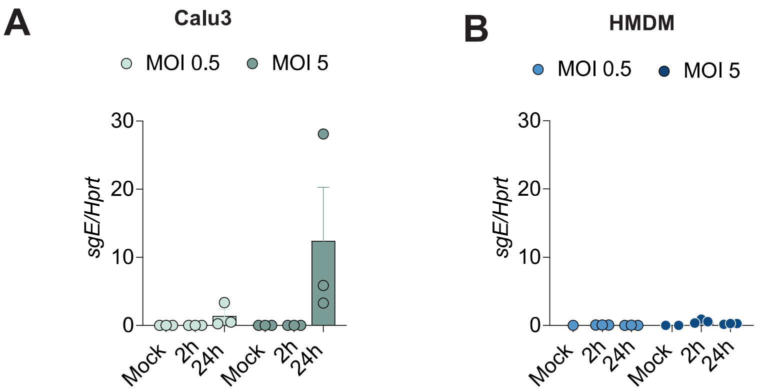


Figure S2: SARS-CoV-2 does not replicate in HMDM

Calu-3 (A) or HMDM (B) were infected as indicated, and the virus was left on the cells. Viral RNA isolated from cells was measured by qPCR. Graphs show mean + SEM, and each point represents an independent experiment (Calu-3; n = 3) or independent donors (HMDM; n = 3).

**Supp Figure 3**

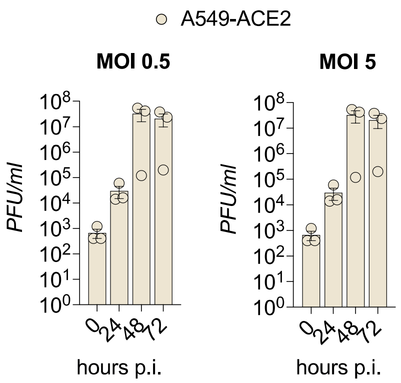


Figure S3: SARS-CoV-2 replicates in ACE2-A549 cells

A549 cells ectopically expressing ACE2 were infected with SARS-CoV-2 at MOI 0.5 or MOI 5. After one h incubation, the virus was removed, and the media was replaced. Cells were harvested at the indicated times (0 h = immediately after the virus inoculum was removed), and viral titers were analyzed by plaque assay.

Supp Figure 4

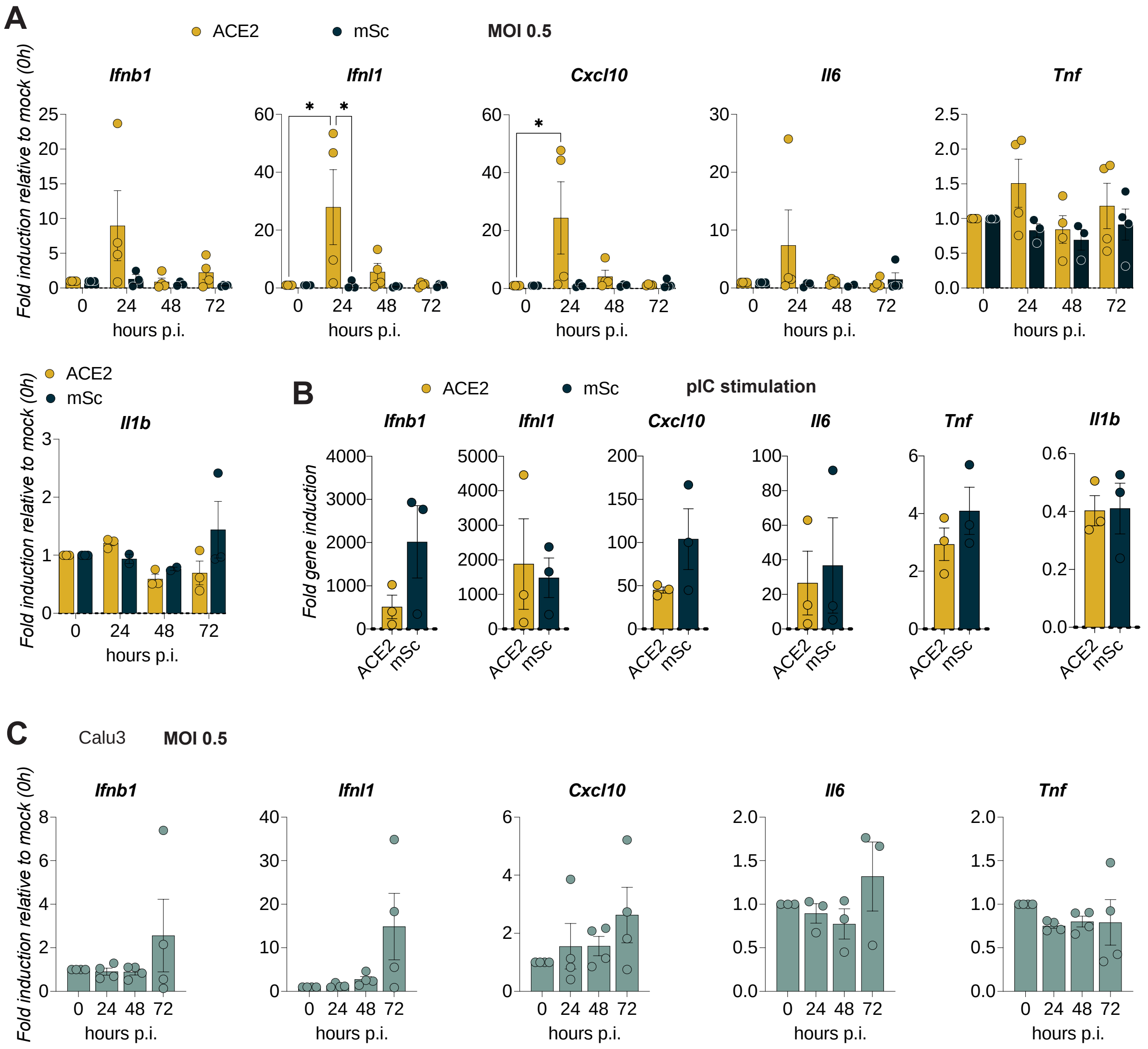


Figure S4: Macrophage ACE2 expression potentiates macrophage inflammatory responses to SARS-CoV-2 at MOI 0.5

A, C: THP-1-ACE-2 , THP-1-mSc (A) or Calu-3 (C) cells were infected with SARS-CoV-2 at MOI 0.5. After one h incubation, the virus was removed, and the media was replaced. Cells were harvested at the indicated times (0 h = immediately after the virus inoculum was removed), and gene expression was quantified by qPCR. Gene expression at each time point is presented relative to the mock control to show fold gene induction. Data show mean + SEM of 4 independent experiments, and significance is indicated by asterisks: $p \leq 0.05$ (*), $p \leq 0.001$ (**), $p \leq 0.0001$ (***) (two-way ANOVA, Tukey's multiple comparison test). (B) : THP-

1-ACE-2 and THP-1-mSc cells were transfected with pI:C for 16 h, and gene expression was assessed by qPCR. Data show fold induction relative to mock-treated cells; graphs are mean + SEM of 4 independent experiments. Significance was assessed using a two-tailed paired t-test.

**Supp Figure 5**

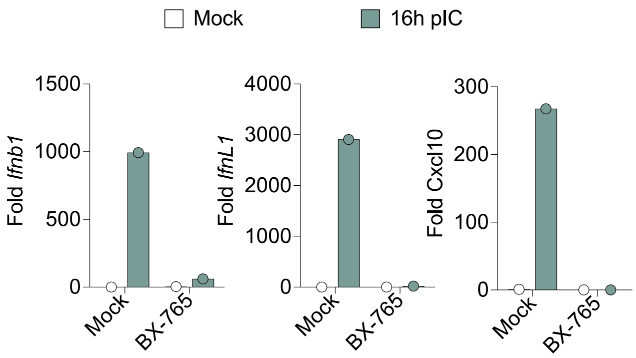


Figure S5: BX-795 blocks anti-viral cytokine induction

THP-1-ACE2 cells were stimulated with pI:C (transfected) for 16 h, with and without BX-765, and gene expression was analyzed by qPCR. Data are mean + SD of duplicate wells and are representative of two independent experiments
